# ClaPNAC: a Classifier of Protein – Nucleic Acid Contacts

**DOI:** 10.1101/2025.05.30.657037

**Authors:** Grigory Nikolaev, Eugene F. Baulin, Janusz M. Bujnicki

## Abstract

Nucleic acid–protein interactions are fundamental to many cellular processes, including genome stability and replication, and the regulation of gene expression. They rely on the formation of specific contacts between amino acid and nucleotide residues. Accurate detection and classification of these inter-residue contacts in both experimentally determined structures and computationally predicted models of nucleic acid–protein complexes are crucial for achieving a comprehensive, molecular-level understanding of nucleic acid–protein recognition mechanisms. Here, we present ClaPNAC, a classifier that annotates 3D structures of nucleic acid–protein complexes with pairwise contacts. It distinguishes stacking interactions (involving the faces of nucleobases), pseudo pairs (involving the Watson-Crick, Hoogsteen, or sugar edge of the base), and phosphate and ribose interactions on the nucleoside/nucleotide side, as well as side chain and backbone interactions on the amino acid side. ClaPNAC is based on a geometric approach, extending our previous method ClaRNA and uses a database of pre-classified doublets extracted from experimentally solved RNA–protein complex structures. Unlike other tools that focus on specific chemical interactions such as hydrogen bonds or van der Waals contacts, ClaPNAC relies on the geometry of the interacting residues. It is applicable to both RNA– and DNA–protein complexes, and it can also operate on isolated nucleosides/nucleotides and amino acids. It can be extended to support additional interaction types and higher-order spatial relationships. ClaPNAC constitutes a notable advancement in the characterization of amino acid–nucleotide interactions by providing a comprehensive classification framework that deepens our insight into these fundamental biological processes. Its applicability to 3D structure prediction and validation highlights its value as a resource for researchers in structural biology.

## INTRODUCTION

Nucleic acid (NA)-protein interactions play essential roles in a wide range of cellular processes, including transcription, translation, splicing, DNA replication, repair, and RNA modification (1–3). Detailed knowledge of these interactions at the molecular level is required to understand the mechanisms of NA-protein recognition and function. Understanding these mechanisms requires structural characterization of NA–protein complexes, as well as methods to analyze and interpret their interaction modes in detail. The idea of a geometric classifier for NA–protein interactions is novel and addresses an unmet need, as current tools either focus on physical interactions (e.g., hydrogen bonds) or lack fine-grained classification. The ability to distinguish pseudo-pairing, stacking, and other contact types based on 3D geometry is particularly useful for annotating predicted or low-resolution models.

NA–protein complexes vary widely in their architecture, dynamics, and stability. Some are highly stable, with precise and conserved contacts, such as ribosomes or CRISPR– Cas effector complexes, where nucleic acids are tightly bound and recognized in a structurally defined manner throughout the complex lifetime (4). Others consist of structured elements that undergo large conformational rearrangements and changes in NA–protein contacts during their functional cycles. A notable example is the spliceosome, a dynamic assembly of small nuclear RNAs and proteins that rearranges multiple times to facilitate intron removal and exon ligation. The interactions between proteins and RNA in this context change both in composition and geometry across different spliceosomal states. Many NA– protein interactions are transient and variable, particularly when involving long non-coding RNAs (lncRNAs) and proteins with intrinsically disordered regions (IDRs). These elements often lack stable tertiary structures in isolation but undergo folding upon binding. Examples include the interaction of Xist lncRNA with heterogeneous nuclear ribonucleoproteins (hnRNPs) during X-chromosome inactivation, and NEAT1 with proteins involved in paraspeckle formation (5). Another well-studied system is the NORAD–PUMILIO complex, where the lncRNA NORAD serves as a multivalent scaffold for PUM proteins through numerous short motifs, relying on both structured and flexible regions (6). Similarly, proteins with disordered domains, such as FUS or TDP-43, can bind RNA in a sequence-independent manner, often contributing to the formation of dynamic membraneless organelles via phase separation (7).

The complex nature of NA–protein interactions has been a subject of intense structural studies, with researchers aiming to elucidate the specific contacts made at the residue level. Structural analyses, initially carried out using X-ray crystallography and nuclear magnetic resonance (NMR) spectroscopy, and more recently using cryo-electron microscopy (cryo-EM), have provided detailed atomic-resolution insights into NA–protein complexes. These studies have revealed key molecular determinants of nucleic acid recognition, as well as recurring interaction patterns observed across both related and unrelated biological systems.

Comparative analyses of experimentally determined NA–protein structures, often in conjunction with sequence conservation data, have shown that binding specificity is frequently mediated by conserved protein domains that recognize distinct nucleic acid sequence or structural motifs. Such domains are commonly found in a broad class of nucleic acid–binding proteins and often act in a modular fashion, combining sequence- and shape-specific recognition features. Extensive structural and functional studies have been performed for both DNA- and RNA-binding proteins, identifying numerous domain families responsible for NA recognition. In DNA-binding proteins, classical motifs such as the helix-turn-helix (HTH), zinc finger, leucine zipper, and homeodomain recognize specific base sequences, often inserting α-helices into the major groove of double-stranded DNA (8). For RNA-binding proteins, domains such as the RNA recognition motif (RRM), K homology (KH) domain, cold-shock domain (CSD), and the double-stranded RNA-binding domain (dsRBD) are commonly involved. These domains show a wide range of structural strategies to accommodate the flexibility and complex folding of RNA molecules, including both sequence- and structure-specific contacts (9).

In general, NA recognition strategies can be broadly divided into two categories: shape readout and base readout. In shape readout, proteins recognize the overall geometry or specific structural motifs of the nucleic acid, such as bulges, hairpins, or grooves. In base readout, the specificity arises from direct interactions with the nucleobases, either through hydrogen bonding or base stacking. Often, both strategies are employed in tandem to ensure tight and specific binding.

Different types of NA-binding domains tend to form distinct types of amino acid– nucleotide contacts at the atomic level. These can include hydrogen bonding to bases or the backbone, stacking interactions between aromatic residues and nucleobases, electrostatic interactions with the phosphate groups, and insertion of structural elements such as helices or loops into nucleic acid grooves. The nature and geometry of these interactions vary depending on whether the target is single- or double-stranded, structured or flexible, and RNA or DNA. For instance, classical RNA-binding domains, such as the RRM or the KH domain, use aromatic side chains (most commonly Phe, Tyr, or Trp) to form stacking interactions with the faces of exposed RNA bases, while additional contacts are made through hydrogen bonds between polar and charged side chains (often Asn, Gln, and Arg) and the edges of the bases or the sugar-phosphate backbone (10). On the other hand, dsRBDs recognize the A-form double helix of RNA mainly through contacts between basic amino acid side chains (in particular Lys and Arg) and the phosphate groups in the RNA backbone. Likewise, DNA-protein interactions rely on specific recognition motifs that mediate base-specific or backbone-based recognition. DNA-binding proteins, such as transcription factors, often use conserved domains like the HTH motif, zinc fingers, and leucine zippers. In zinc finger domains, sequence specificity arises from hydrogen bonds between side chains (typically from R, H, N, and Q) and the edges of DNA bases exposed in the major groove (11). The basic region-leucine zipper (bZIP) domain mediates sequence-specific DNA binding through a basic α-helix inserted into the major groove, where charged residues (Lys and Arg) form hydrogen bonds and electrostatic interactions with base edges and phosphate groups (12). Some proteins interact with DNA in a sequence-independent manner, particularly when recognizing the shape and electrostatics of B-form DNA. Examples include histones in nucleosomes and architectural proteins like HMG-box or HU/IHF. They often interact primarily with the DNA phosphate backbone through basic residues (Lys and Arg) and insert minor-groove-binding motifs, such as AT-hooks or intercalating residues (typically Arg or Pro), that contact nucleobases nonspecifically or distort DNA conformation.

As experimentally determined structures of NA–protein complexes accumulated, various studies began to statistically analyze pairwise contacts between nucleotide and amino acid residues, aiming to uncover general preferences that drive molecular recognition. One of the earliest efforts focused on DNA–protein interactions, with researchers using large datasets of crystal structures to identify common patterns of amino acid–base contacts. These studies revealed the frequent involvement of basic residues such as arginine and lysine in contacting the DNA backbone, as well as the recurring use of α-helices inserted into the major groove to facilitate sequence-specific recognition (13). In a focused analysis of RNA–protein interfaces, Treger and Westhof statistically surveyed 45 crystal structures of RNA–protein complexes (14). Their findings highlighted the dominance of certain amino acids, particularly Arg and Lys, in interacting with the RNA phosphate backbone. Although they found no strong preference for specific RNA bases at the interface, residues such as Pro and Asn showed mild preferences for interacting with particular bases. Another important insight from this and related work was the frequent involvement of water molecules in bridging protein and RNA components, mediating interactions that could not occur through direct contacts alone (13). Another study explored the recognition of nucleic acid bases by amino acid side chains through hydrogen bonding (15). Their analysis revealed that only a few amino acids, including Asn, Gln, and Arg, could form specific hydrogen bonds with nucleotide bases in a coplanar fashion. Such direct interactions involving specific hydrogen bonds are the key for sequence-specific recognition.

Kondo and Westhof explored how nucleotide bases interact with amino acid residues across a variety of DNA/RNA-binding and nucleotide-binding proteins (16). They analyzed 428 crystal structures of nucleotide–protein complexes, encompassing both full nucleic acids and isolated nucleotides or nucleosides. The interactions observed between nucleobases and amino acids were then categorized based on their hydrogen-bonding patterns. As a result, they introduced the concept of “pseudo pairs”—that is, configurations where amino acid residues form hydrogen bonds with nucleotide bases, mimicking the base pairing seen in DNA and RNA structures. The authors identified that only five out of the twenty standard amino acids, namely Asn, Gln, Asp, Glu, and Arg, possess the necessary planar structures with hydrogen-bonding donor and/or acceptor atoms that resemble those in nucleobases. Additionally, they found that the peptide backbone of proteins has the potential to form pseudo pairs, particularly showing a strong preference for binding to the adenine base.

While the study of Kondo and Westhof provided a classification scheme to characterize certain classes of nucleotide-amino acid interactions, thus far no computational program was developed to analyze structures of RNA-protein, DNA-protein or nucleotide/nucleoside-protein complexes and generate annotations that distinguish between different types of interactions. Such programs have been developed for contacts within nucleic acids, e.g., RNAView (17), MC-Annotate (18), FR3D (19), DSSR (20), and ClaRNA (21), and they can be used to identify specific base pairs, stacking interactions, base-phosphate contacts and other pairwise interactions. To enable similar analyses of NA-protein interactions, we developed a classifier for amino acid-nucleotide/nucleoside interactions ClaPNAC, which takes as input 3D structures and annotates them with pairwise contacts that distinguish stacking interactions (involving the faces of nucleobases), pseudo pairs (involving the Watson-Crick, Hoogsteen, or sugar edge of the base), and phosphate and ribose interactions on the nucleotide/nucleoside side, and side chain and backbone interactions on the amino acid side. ClaPNAC can detect specific types of interactions that are prevalent in nucleic acid/nucleotide/nucleoside-protein structures but often overlooked by other methods. This method is designed to easily incorporate additional interaction types in the future. It can also be used to identify recurrent structural motifs.

## MATERIALS AND METHODS

### Classification scheme and data preparation

The stages of developing ClaPNAC are presented in **Figure 1**. We used the in-house non-redundant dataset of 2,164 high-resolution RNA-protein complexes from the PDB (22) (release date: October 14, 2022). A randomly selected subset of 100 structures was used as a test dataset, while the remaining structures were used for training. The lists of structures included in the training and test datasets are provided in **Supplementary File S1**. For all structures, we identified nucleotide-amino acid doublets that were close in space, defined by having at least two pairs of non-hydrogen atoms within a distance of 3.5 Å, that is the average distance for weak hydrogen bond (23). Each doublet is uniquely described by its PDB ID, chain ID, residue type, and the numbers of the residues and chains involved. For example, ADE_ALA_1JJ2_0_1328_X_166 denotes a doublet from PDB file 1JJ2, with the first residue being adenosine number 1328 in chain 0, and the second residue being alanine number 166 in chain X. All doublets were divided into subgroups by the involved residues (**Supplementary Table 1**). We applied the clustering methods HDBSCAN (24) and SimRNAweb 2.0, Option C (25) to the subgroups of doublets superimposed by the nucleotide. The clustering shows the most preferable type of contact for each subgroup (**Figure 2A**). We declared seven interaction classes: 1) amino acid contact with phosphate group of the nucleotide (class P); 2) contact with ribose (class R); 3) with Hoogsteen edge (class H); 4) with sugar edge (class S); 5) with Watson-Crick edge (class W); 6) stacking with + face of nucleotide base (class +); 7) stacking with - face of nucleotide base (class -) (**Figure SF1, Figure 2B**).

**Figure 1.**
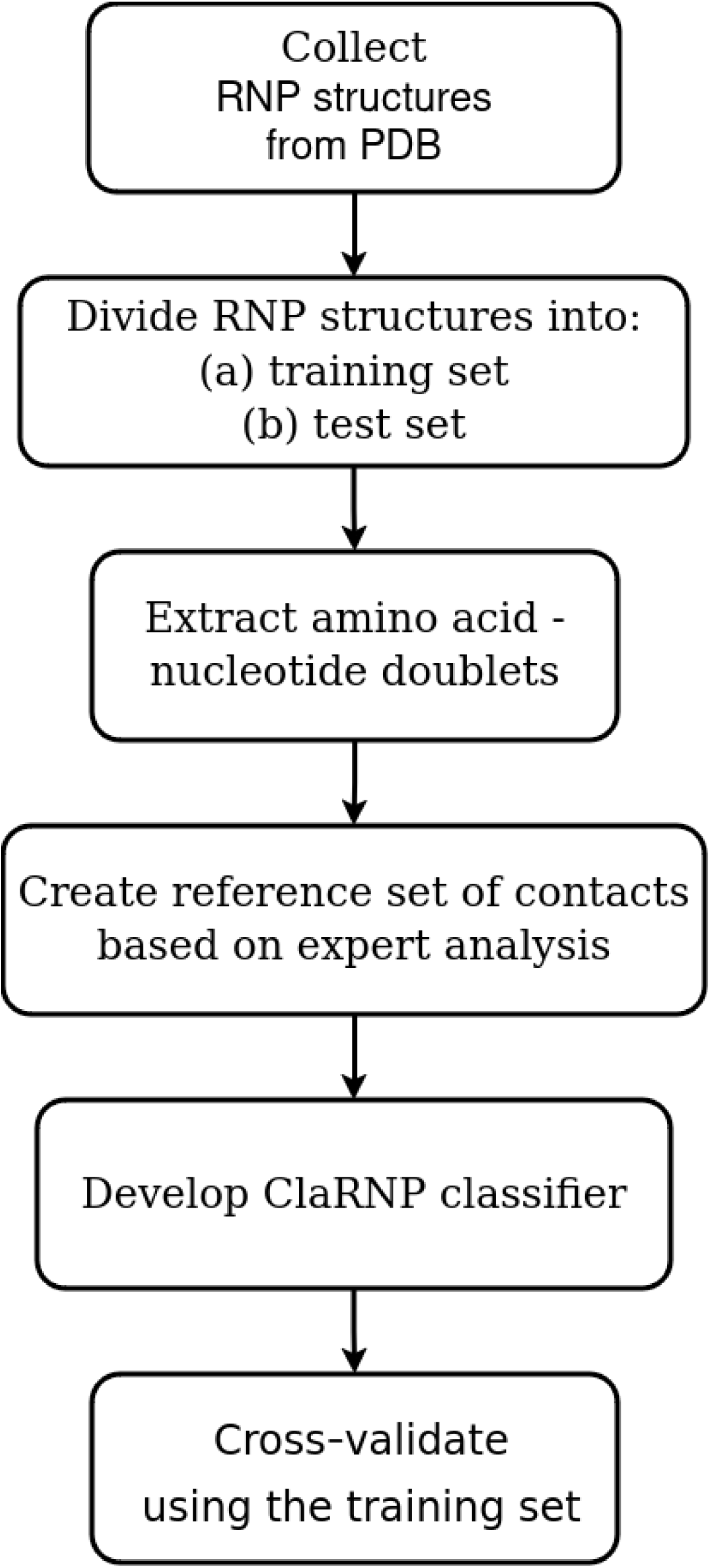
Pipeline for training ClaPNAC on structures from the PDB database.

**Figure 2.**
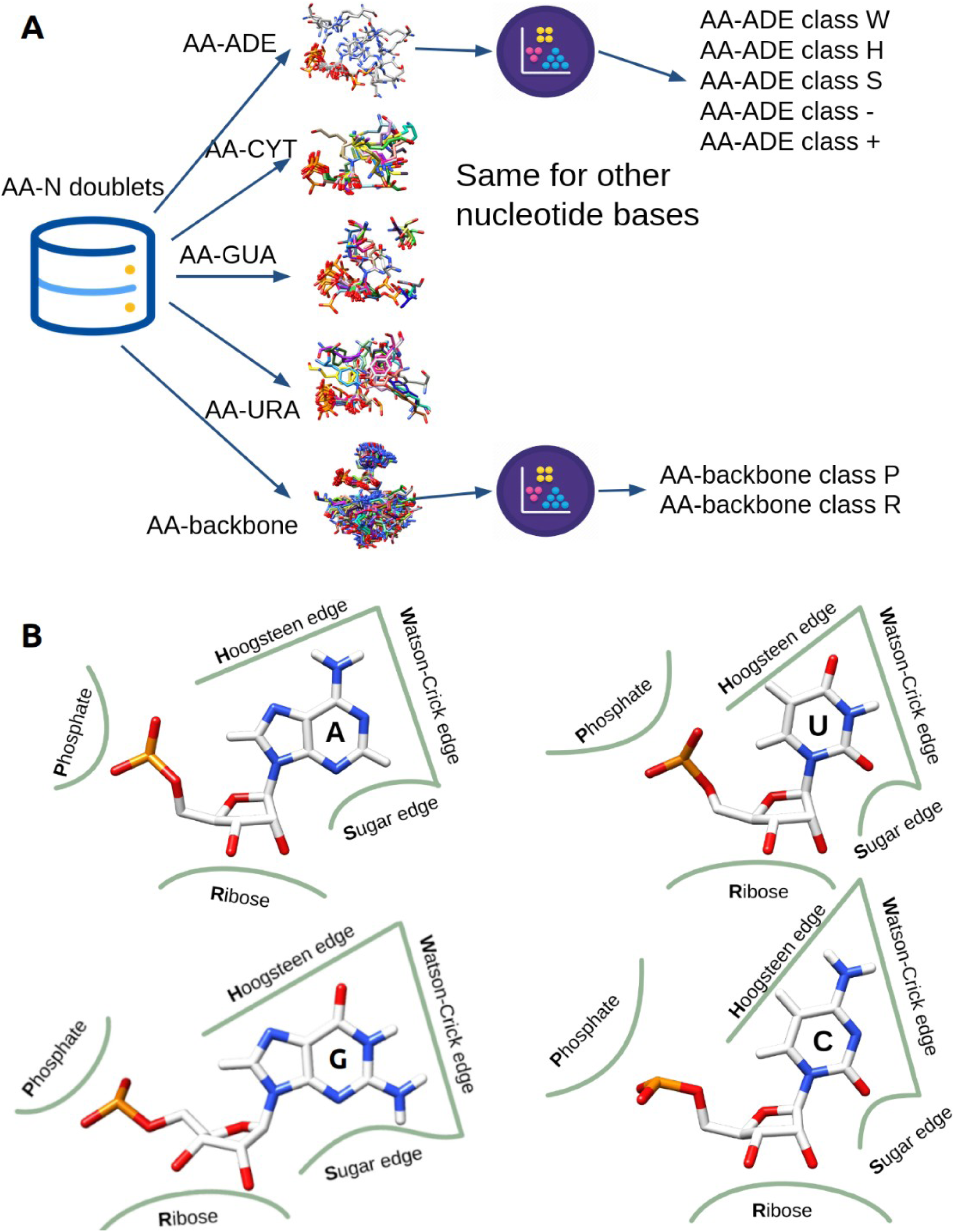
Training dataset preparation. (**A**) Pipeline for extracting representatives of each interaction class for ClaPNAC. (**B**) ClaPNAC interaction classes for adenine, guanine, uracil, and cytosine are in line with the nomenclature suggested in (16). The regions where amino acid residues form pairs with nucleotides, and are labeled with a specific class, are indicated by green lines.

Before finalizing the training set, we conducted cross-validation on all members across all classes. For each class (W, H, S, +, -, P, or R), a doublet was selected. ClaPNAC was then trained while excluding this doublet, and the model’s ability to correctly classify the excluded doublet as a representative of its specific class was evaluated.

### ClaPNAC algorithm

The ClaPNAC algorithm is based on a geometric approach and allows for the annotation of amino acid – nucleotide doublets according to the classes described above. The first step is to extract nucleotide-amino acid doublets from the input structure (i), then superimpose each doublet by nucleotide base or backbone with the representatives of the possible classes (ii). For example, the doublet ADE_ALA_1JJ2_0_1328_X_166 will be superimposed by the adenine base for W, H, and S classes, and by the backbone for P and R classes. The next step is to calculate the **d^1^** = RMSD values between the atoms of amino acids of each class member i (from 1 to n) and the atoms of the amino acid from the input doublet ((n+1)th doublet), as well as the **d^2^** = dRMSD (RMSD of pairwise atom distances between the amino acid and the nucleotide base/backbone) values (iii). Finally, the probability of belonging to a certain interaction class is estimated with the scoring function SF (Figure 3**, Formula 1,2**) (iv).

**Figure 3.**
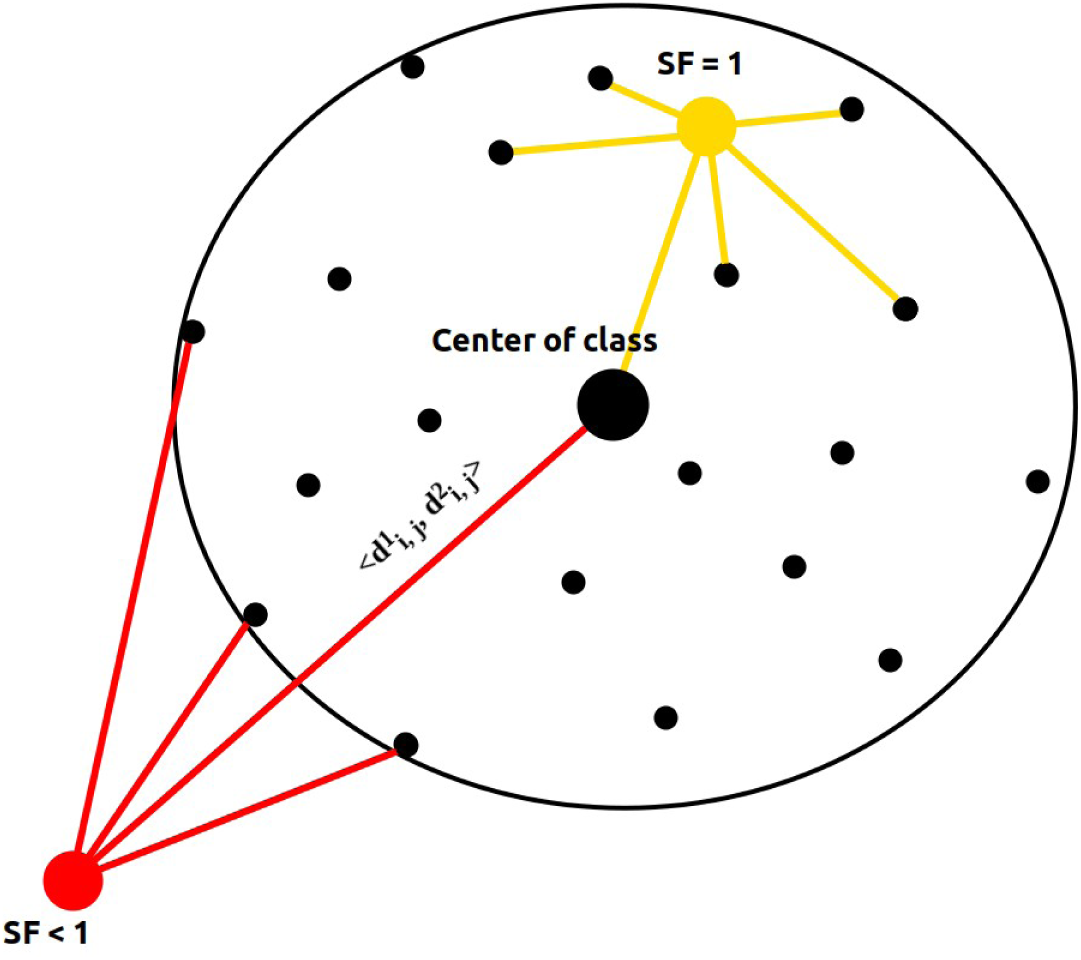
ClaPNAC scoring function

The behavior of the ClaPNAC scoring function is illustrated using black dots to represent class members, a large black dot to indicate the class center, and yellow and red dots to represent input doublets that are being evaluated for potential class membership.

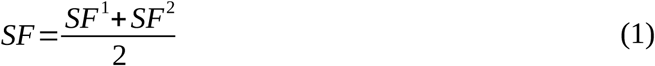

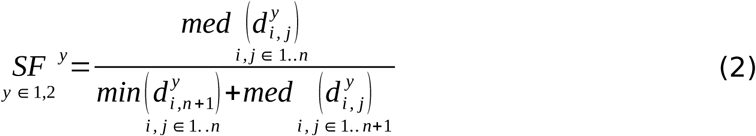

The ClaPNAC scoring function enables us to assess the position of the amino acid from the input doublet, in comparison to the amino acids of the class members, when nucleotides are superimposed by base or backbone. If SF is more than 0.5, it indicates that the input amino acid is closer to the center of the class than some other members. If SF is less than 0.5, the input amino acid is “near” the class, meaning it is farther from the center, resulting in a lower SF score. For each interaction class, we manually defined the threshold for the SF, based on the density of class representatives in the training dataset. For example, for the ARG class P, the threshold is 0.63, meaning that SF > 0.63 is converted to SF = 1, and SF < 0.63 is normalized by 0.63.

ClaPNAC is also capable of classifying nucleotide–amino acid pairs within a coarse-grained representation. In this mode, nucleotides are represented using five key atoms—P, C4′, N9, C2, and C6 for adenine and guanine, and P, C4′, N1, C2, and C4 for cytosine and uracil. Amino acids are represented using 2 to 4 atoms, such as CB, NE, NH1, and NH2 for arginine, or CB and SG for cysteine, among others.

### Benchmarking

As no established benchmarks or comparable tools exist for this specific type of classification, the assessment relied on indirectly comparable tools as follows. We used DSSR version 2.0 (26) and fingeRNAt (27) to define hydrogen bonds, van der Waals interactions and stacking for the input doublets, and then assigned the interaction classes according to the following rules:

1. If amino acid atoms interact with nucleotide atoms P, OP1 or OP2, the interaction is categorized as class P.
2. Stacking interactions are classified as “stacking” without separation into (+) and (−) classes.
3. If ribose atoms O5’, C5’, C4’, O4’, C3’, O3’, C2’, O2’ are involved, then interaction is recognized as class R.
4. For specific base types, the classifications are as follows:

### Adenine

#### Guanine

Class H: Atoms C8, N7, C5, N6.

Class W: Atoms C2, N1.

Class S: Atoms N3, C4.

Class H: Atoms C8, N7, C5, O6.

Class W: Atoms N1, N2.

Class S: Atoms N9, N3.

#### Cytosine

Class H: Atoms C5, N4, C4.

Class W: Atoms N3.

Class S: Atoms O2.

#### Uracil

Class H: Atoms C5, O4.

Class W: Atoms N3.

Class S: Atoms N1, O2.

This classification scheme allows for the systematic identification of interactions based on the specific atomic components involved. Using the above definitions, ClaPNAC has been benchmarked using the test dataset, which includes 100 randomly selected RNA-protein complexes, which was not used in the training dataset.

### Software

The new NA-protein contact classifier (ClaPNAC) and additional scripts, including a contact extractor utility (ContExt), were developed based on the Python programming language, as well as Biopython (28) and SciPy (29) libraries.

### Hardware

For calculations with third-party methods and for the training and benchmarking of the ClaPNAC classifier, we used an in-house high-performance computing cluster comprising 624 nodes (each with a 2.2 GHz processor and 2 GB of RAM).

## RESULTS

### Comparison with other tools

ClaPNAC is a multi-class, multi-label classifier, as each doublet can be annotated with multiple interactions. Therefore, its evaluation is more complex than that of classical binary classifiers.

We compared ClaPNAC with DSSR 2.0 and fingeRNAt following the rules described in the Methods section (Figure 4). DSSR identified 1,605 doublets, fingeRNAt identified 2,168, and ClaPNAC labeled 1,379 doublets with a score of 1.0. Assuming the DSSR results as the true positive set, the resulting recall score for ClaPNAC was 0.29. All three tools classified 405 doublets (P – 259, R – 96, W – 14, H – 15, S – 21, stacking – 0), with 87% of these doublets involving interactions with the nucleotide backbone and only 13% involving bases. The limited number of interactions with bases can be explained by differences in approach among the tools (geometric for ClaPNAC versus real binders for DSSR and fingeRNAt) and by the strict criteria for selecting training classes for ClaPNAC (for classes W, H, and S, the nucleotide base and amino acid side chain must be coplanar and contain at least two pairs of non-hydrogen atoms within a distance of 3.5 Å).

**Figure 4.**
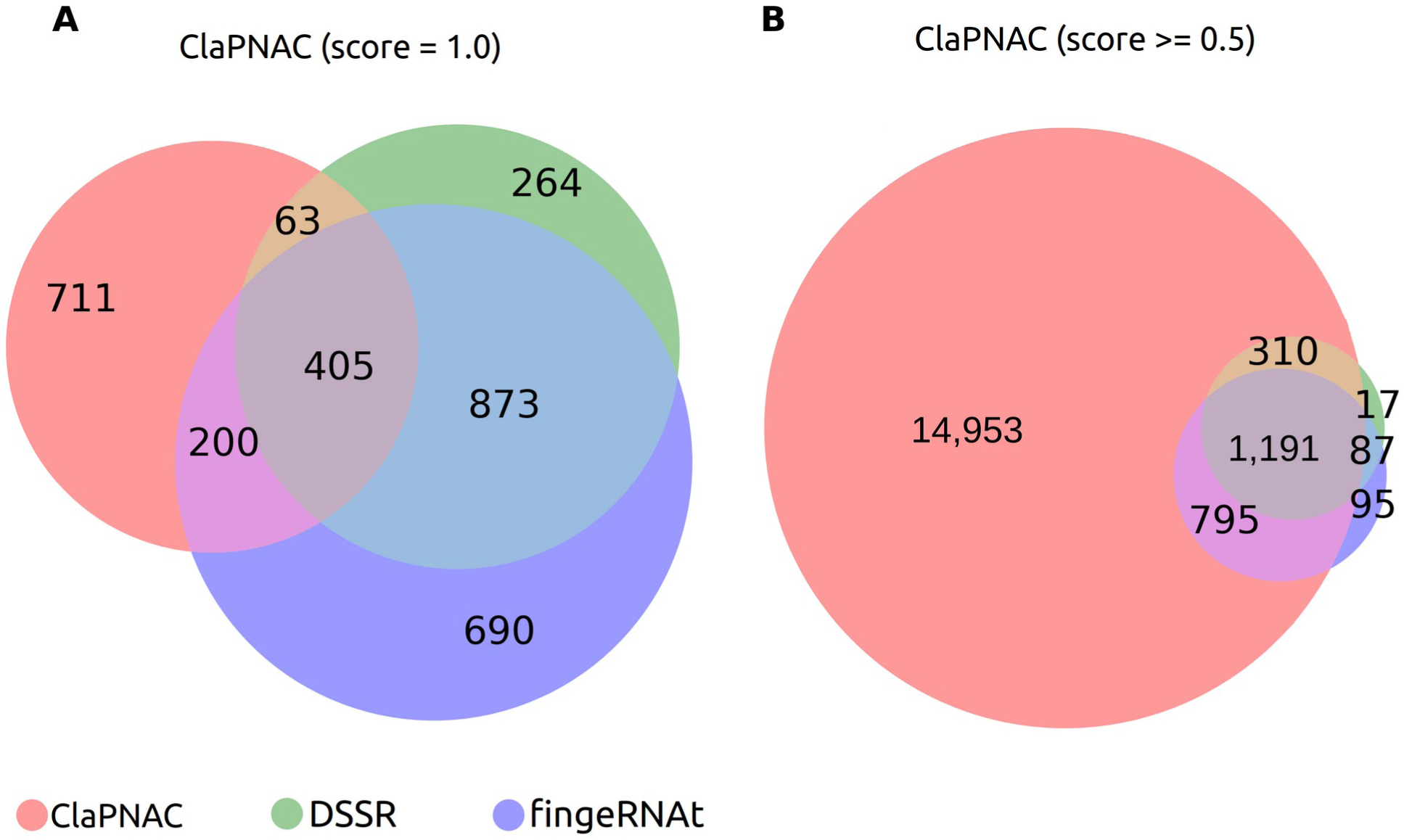
Comparison of the ClaPNAC, DSSR and fingeRNAt results for the test set. The following results were obtained: **(A)** With SF = 1.0 ClaPNAC classified 1,379 doublets, fingeRNAt – 2,168, DSSR – 1,605. ClaPNAC identified 711 doublets, that were not classified by other tools, fingeRNAt – 690 and DSSR – 264. **(B)** With SF ≥ 0.5 ClaPNAC classified 17,249 doublets and missed only 104 doublets identified by DSSR, and 182 doublets identified by fingeRNAt.

ClaPNAC labeled 17,249 doublets with a score greater than 0.5 and failed to classify only 199 doublets that were classified by DSSR or fingeRNAt. Assuming the DSSR results as the true positive set, the resulting recall score for ClaPNAC was 0.94. Most of the cases that ClaPNAC failed to classify involve only one hydrogen bond, and all other amino acid atoms are more than 3.5 Å away from any nucleotide atom. They are also located far from the “cloud” of training doublets (**Figure SF2**).

Notably, fingeRNAt and DSSR detect contacts at a distance of 6.0 Å, and in the ClaPNAC training dataset there are doublets only with at least two pairs of non-hydrogen atoms within a distance of 3.5 Å, so doublets with a larger interaction distance between nucleotide and amino acid residue will have lower score. Figure 4B shows that at a lower threshold (≥ 0.5) ClaPNAC can cover approximately 93% of doublets found by DSSR and fingeRNAt. We benchmarked the time required to process the PDB structures from the dataset with 100 randomly selected structures. ClaPNAC is three times slower than DSSR, but more than 30 times faster than fingeRNAt. To analyze the test dataset ClaPNAC required 1m 29s, DSSR – 36s and fingeRNAt – 20m 48s.

### Benchmarking on the labeled dataset

The dataset from the study by Kondo and Westhof (16) was also used as a test dataset. The lists of structures included in the training and test sets are provided in Supplementary File ***S1***.

Kondo and Westhof found solid representatives for W, H and S classes in the dataset with 428 structures, that includes nucleotides and amino acids. The results of classifying this dataset with ClaPNAC as shown in Figure 5A and **5B**. With score = 1.0 ClaPNAC can correctly classify 76% of the doublets. Missing doublets in most cases could be divided into two groups: 1) doublet makes hydrogen bond with additional O atom from water molecule (e.g., PDB id: 2OXC, ADE_ARG_2OXC_A_300_A_84, class W); 2) residue and nucleotide are not in one plane (e.g., PDB id: 1JJV, ADE_ASN_1JJV_A_300_A_175, class H). ClaPNAC identified 79 additional doublets (Figure 5C**, 5D, 5E**). Each of these doublets includes two independent hydrogen bonds involving four distinct atoms and represents W classes not identified by Kondo and Westhof (16). These additional doublets were detected due to the ClaPNAC geometric approach, which relies on the spatial orientation of molecules in 3D space rather than explicit hydrogen bond detection (Kondo and Westhof recognize the doublet, if it has at least two hydrogen bonds). With a score ≥ 0.69, ClaPNAC correctly classified all the doublets identified by Kondo and Westhof and identified 174 additional doublets.

**Figure 5.**
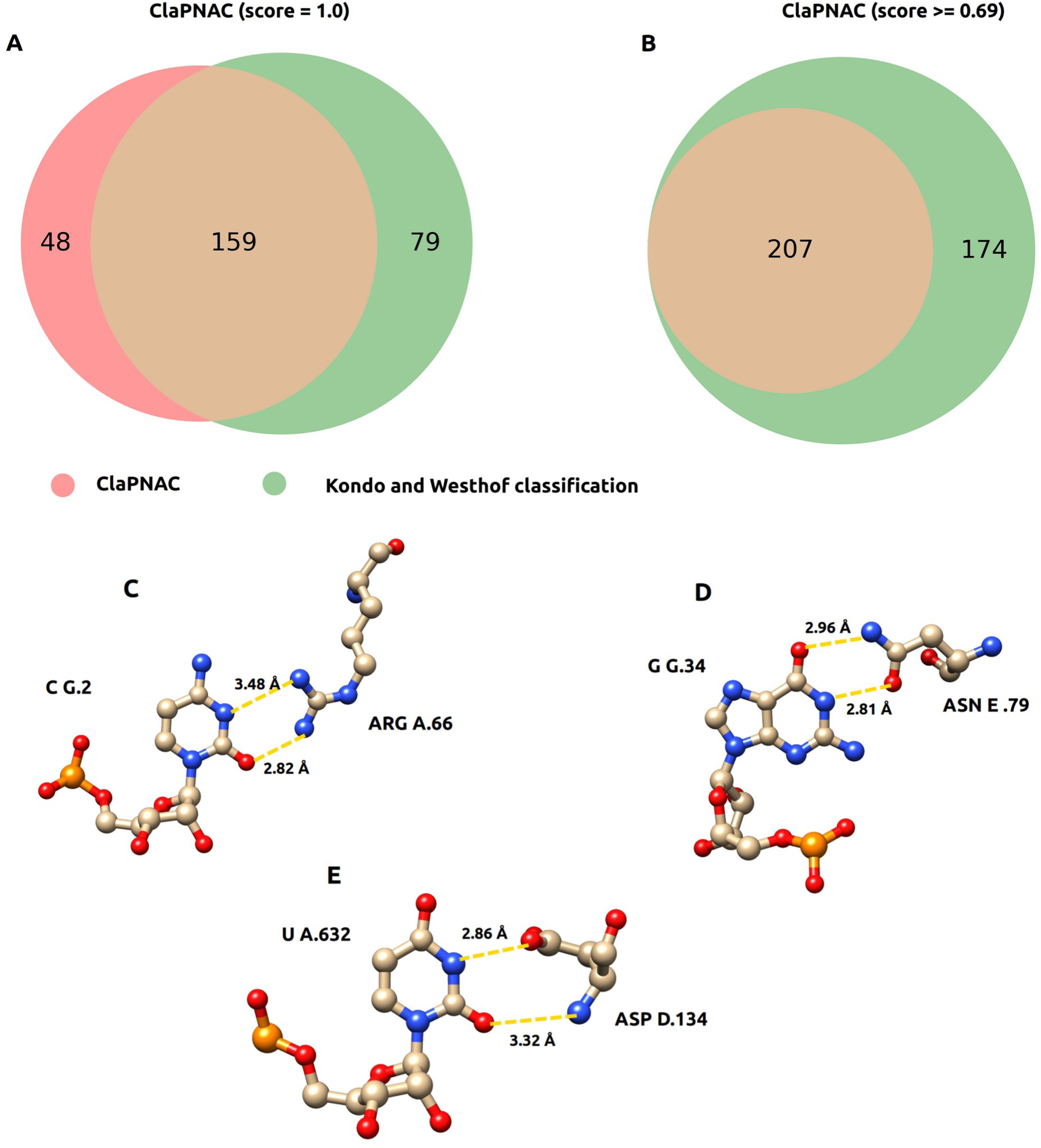
ClaPNAC results for the Kondo and Westhof dataset. The following results were obtained: **(A)** with the score = 1.0 ClaPNAC classified 238 doublets in total, including 159 doublets that were labeled by Kondo and Westhof**. (B)** With the score ≥ 0.69 ClaPNAC classified 381 doublets in total, including all 207 doublets that were labeled by Kondo and Westhof. Examples of doublets that were not labeled by Kondo and Westhof but classified by ClaPNAC with the score = 1. **(C)** Interaction class W for C2 chain G and ARG66 chain A from structure PDB id: 1PVO (30), **(D)** interaction class W for G34 chain G and ASN79 chain E from structure PDB id: 2XDD (31), **(E)** interaction class W for U632 chain A and ASP134 chain D from structure PDB id: 5IWA (32).

No structures from Kondo and Westhof dataset were used for training ClaPNAC.

### Use case 1: Contact analysis for U5 snRNP State I, II, III, IV and U4/U6.U5 tri-snRNP

The human U5 snRNP is the ∼1 megadalton “heart” of the spliceosome and is recycled through an unknown mechanism involving major architectural rearrangements and the dedicated chaperones CD2BP2 and TSSC4 (33). Using ClaPNAC, we identified differences in the interactions between U5 snRNA and the pre-mRNA-processing splicing factor 8 (PRP8) in complexes 8Q7V (State I), 8Q7Q (State II), 8Q7W (State III), and 8Q7X (State IV) (33), as well as 6QW6 (U5.U4/U6 tri-snRNP) (34). ClaPNAC reveals differences in the conformation of U5 snRNA and PRP8 (**Supplementary Table 2,** Figure 6).

**Figure 6.**
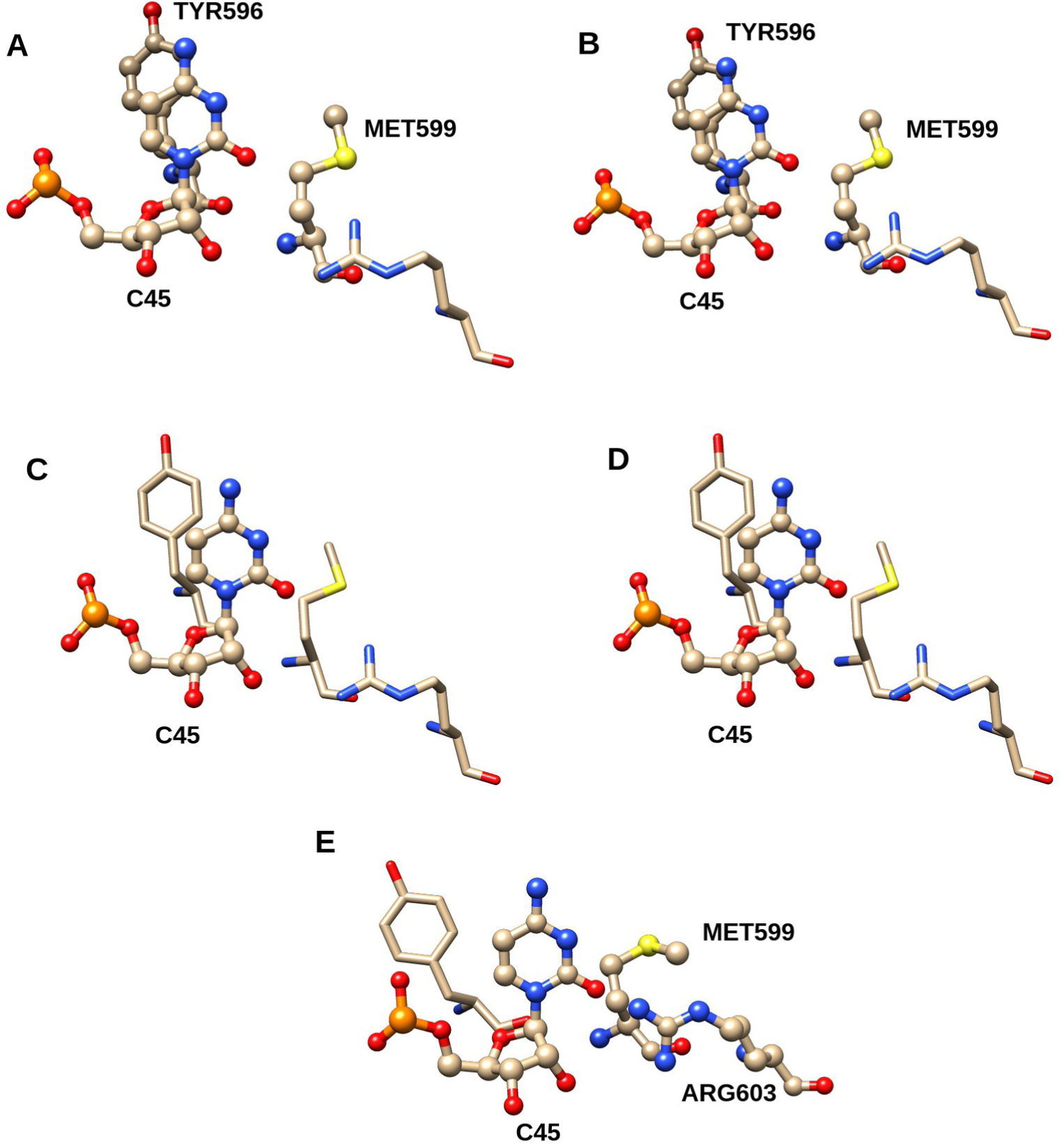
Examples of the similarities and differences in the structures of U5 snRNA and pre-mRNA-processing splicing factor 8 in complexes 8Q7V, State I (A); 8Q7Q, State II (B); 8Q7W, State III (C); 8Q7X, State IV (D); and 6QW6, U5.U4/U6 tri-snRNP (E). Amino acid residues with ClaPNAC interaction scores of 1.0 are labeled and shown in the ball-and-stick model, while residues that do not form interactions are unlabeled and shown in the stick model.

States I and II have the same interaction patterns, as do States III and IV, as shown in Figure 6 and **Supplementary Table S2**. However, there is a significant difference between States I/II and States III/IV (33). Interestingly, the U5.U4/U6 tri-snRNP exhibits some contacts similar to those in States I/II (e.g., G20–Pro463), others resembling those in States III/IV (e.g., C13–Thr224), and unique contacts not found in either group (e.g., U17–Arg470). This type of analysis is not readily achievable with existing tools such as DSSR and fingeRNAt.

### Use case 2: Sensitivity to small changes

To demonstrate sensitivity of ClaPNAC to small structural differences, we analyzed three highly similar RNP complexes of the c-di-GMP riboswitch bound to c-di-GMP with small nuclear ribonucleoprotein A (PDB IDs: 3MUM, 3MUR (35), and 3IRW (36)). The RMSD between these structures is less than 1 Å, yet ClaPNAC was able to detect differences, as shown in **Supplementary Table S3** and Figure 7.

**Figure 7.**
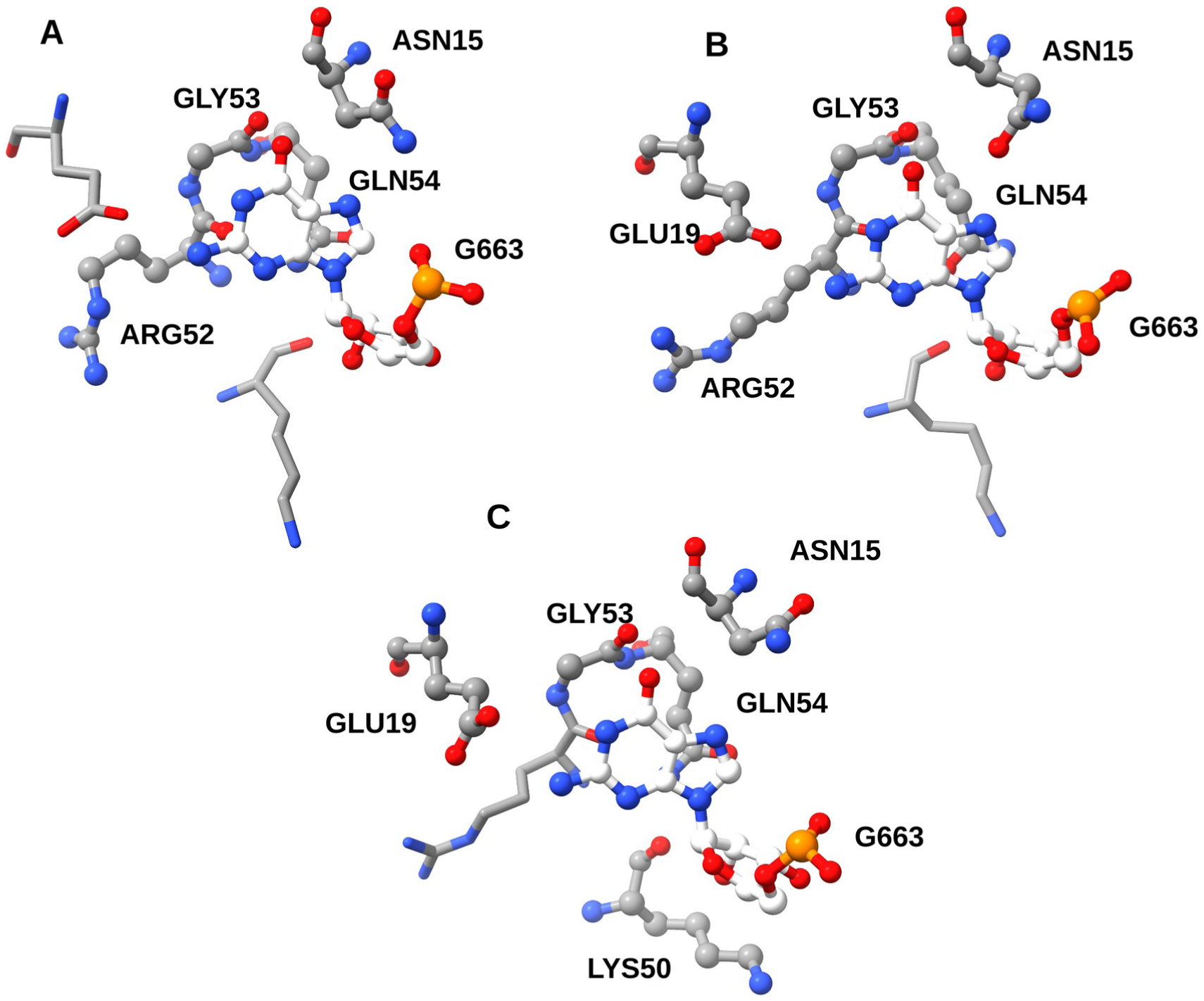
Comparison of the ClaPNAC results with score = 1.0 for the doublets with G663 for structures (**A**) 3IRW, (**B**) 3MUM, (**C**) 3MUR.

Notably, most differences in doublet classification occur in the non-base classes P (e.g., A62-Lys22), R (e.g., G663-Lys50), and S (e.g., U661-Arg52), and in some cases differences for classes + (e.g., C664-Lys88) and - (e.g., A665-Phe56). Figure 7 illustrates the differences in doublet classification involving nucleotide G663 from loop II of the c-di-GMP riboswitch.

### Use case 3: Analysis of Pfam families

We used ClaPNAC to analyze amino acid–nucleotide contacts in two RNA recognition motifs, PF00076 (RRM_1) and PF04059 (RRM_2), and compared the statistics of preferred nucleotide base or backbone interactions, interaction classes, and the nucleotides and amino acids involved (**Supplementary Table S4**). The analysis detected significant differences between the two domains: 1) In the PF00076 domain, RNA predominantly forms contacts with the protein from the backbone side (class P), whereas in PF04059, the preference is reversed (class W); 2) The PF00076 domain shows no nucleotide preference in interactions, while in PF04059, uracil (U) is the most common; 3) In PF00076, residues arginine (Arg) and lysine (Lys) are the most frequently involved in classified doublets, but no such preferences are observed in PF04059. These findings provide a deeper understanding of the specific features of each domain.

### Use case 4: Comparative analysis of RNA-protein and DNA-protein complexes

We used ClaPNAC to analyze amino acid–nucleotide contacts in two datasets: RNA– Protein (100 complexes) and DNA–Protein (47 complexes) (37), then compared statistics of preferred nucleotide base or backbone interactions, interaction classes, and involved nucleotides and amino acids (Supplementary Table S4). The analysis revealed differences between the two datasets: 1) In the DNA–protein dataset, doublets with class P interactions occur more frequently than in the RNA–protein dataset, even though the frequencies of base and backbone interactions are very similar between the two datasets; 2) In the DNA–protein dataset, class H interactions are much more common than class W or class S interactions compared to the RNA–protein dataset.

The first difference can be explained by the additional O2′ oxygen atom in ribonucleotides compared to deoxyribonucleotides. The second difference is explained by the fact that DNA more frequently forms double-stranded tertiary structures, whereas RNA often has single-stranded regions (38), and because in the B-form helix (DNA), the major groove (H edge) is more accessible than in the A-form helix (RNA). Consequently, nucleotides in DNA are more likely to form class H interactions with other nucleotides when Hoogsteen edges are accessible for contacts. The lower frequency of class S interactions in DNA is also linked to the lack of the O2′ oxygen atom. ClaPNAC demonstrates its capability to detect these differences, highlighting its suitability for analyzing both RNA- and DNA-protein structures.

## DISCUSSION

In this work, we present ClaPNAC, a classifier designed to analyze amino acid– nucleotide/nucleoside interactions. Taking 3D structures as input and annotating them with pairwise contacts, ClaPNAC distinguishes various interaction types, including stacking interactions involving the faces of nucleobases; pseudo pairs engaging the Watson-Crick, Hoogsteen, or sugar edges of the base; and interactions with the phosphate and ribose moieties on the nucleotide/nucleoside side, alongside interactions with the side chain and backbone of the amino acid.

The geometric approach employed by ClaPNAC optimizes the classification process by bypassing the problems associated with identifying hydrogen bonds (hydrogen bond criteria vary, e.g., distance and angle thresholds, leading to inconsistencies in identification) and van der Waals interactions (interaction strength depends on atomic radii and well-depth parameters, which vary across force fields, e.g., CHARMM, AMBER, GROMOS). This methodology emphasizes the distinction between common pseudo pairs and interactions involving phosphates and riboses, extending our understanding of the geometry of interaction sites between proteins and RNA. Moreover, this approach enables the systematic identification of similar binding sites in ribonucleoprotein (RNP) structures, which is crucial for deciphering the underlying mechanisms of RNA-protein interactions.

ClaPNAC demonstrates both strengths and limitations in detecting and annotating doublets. While it labeled fewer doublets with a perfect score of 1.0 (1,379 compared to 1,581 by DSSR and 2,131 by fingeRNAt), its geometric approach enabled the identification of unique interactions, such as doublets characterized by strict co-planarity and specific distances between nucleotide bases and amino acid side chains. Moreover, ClaPNAC’s ability to choose a score threshold (e.g., ≥ 0.5) allows it to capture approximately 93% of the doublets identified by DSSR and fingeRNAt, showing its broad applicability. Differences in detection can be attributed to methodological variations. While DSSR and fingeRNAt use larger distance thresholds (up to 6.0 Å), training dataset of ClaPNAC is based on strict criteria requiring at least two pairs of non-hydrogen atoms within 3.5 Å. Thus, ClaPNAC tends to exclude doublets with weak interactions, ensuring higher geometric specificity. Although ClaPNAC is slower than DSSR, its runtime (1m 29s, for 100 structures) is more efficient than fingeRNAt, making it a practical choice for large-scale analyses.

When benchmarked with the dataset by Kondo and Westhof (2011), ClaPNAC demonstrated a high level of classification, correctly annotating 76% of doublets with a score of 1.0. The unclassified doublets were primarily due to differences in classification criteria: ClaPNAC missed interactions requiring additional hydrogen bonds or lacking co-planarity. Nevertheless, ClaPNAC identified 79 additional doublets not detected by Kondo and Westhof, highlighting its ability to uncover interactions beyond conventional hydrogen-bond criteria. By lowering the score threshold to 0.69, ClaPNAC successfully classified all doublets identified by Kondo and Westhof, along with 174 additional interactions, demonstrating its flexibility and precision.

ClaPNAC identified variations in nucleotide-protein interactions across different structural states of U5 snRNP. By detecting unique contacts within the U5.U4/U6 tri-snRNP complex (e.g., U17-Arg470), ClaPNAC revealed the architectural rearrangements that occur during the spliceosomal cycle. These results underscore ClaPNAC’s potential for studying the structural mechanisms that drive complex biological processes.

ClaPNAC also detected subtle differences among highly similar structures of the c-di-GMP riboswitch bound to c-di-GMP. Despite an RMSD of less than 1 Å between these structures, ClaPNAC identified distinct interactions between nucleotides and amino acids, highlighting its precision in capturing small variations in nucleic acid–protein complexes.

The comparison of RNA recognition motifs (PF00076 and PF04059) revealed notable differences in nucleotide interaction classes, highlighting domain-specific features. For example, the PF04059 domain showed a preference for uracil (class W), whereas PF00076 exhibited no nucleotide bias and predominantly interacted with RNA backbones (class P). These findings align with the known functional distinctions between these domains, demonstrating capability of ClaPNAC to capture detailed structural differences within RNA-protein families.

ClaPNAC effectively identified key differences between RNA-protein and DNA-protein complexes. For instance, class P interactions occurred more frequently in DNA-protein complexes, while class H interactions were more common due to the greater accessibility of the major groove in the B-form DNA helix. In contrast, the additional oxygen atom in ribonucleotides contributed to unique interaction patterns in RNA-protein complexes. These findings highlight capability of ClaPNAC to accurately analyze both RNA and DNA interactions.

Moreover, ClaPNAC supports a coarse-grained mode that enables classification of doublets even when some atoms are missing in the input structure. While this mode may reduce accuracy, it increases utility of ClaPNAC for analyzing incomplete or lower-resolution structural data, making it a flexible tool for researchers.

The insights gained from classification of amino acid–nucleotide interactions made by ClaPNAC can greatly contribute to advancing methods for 3D structure prediction and evaluation. By providing a clear framework for understanding interaction geometries, ClaPNAC can help refine existing computational models used to predict the 3D conformations of RNA–protein and DNA–protein complexes.

ClaPNAC has certain limitations: 1) For accurate classification, all residues and nucleotides in the input structure must have all their atoms. If some doublets in the input structure have missing atoms, ClaPNAC can operate in coarse-grained mode, but this results in reduced accuracy. 2) Nucleotides should not be modified; if modified nucleotides are renamed to standard forms, any additional atoms or groups will be ignored.

The ability to classify interactions into specific categories enables more accurate training of machine learning algorithms for structure prediction. For instance, incorporating ClaPNAC interaction data into existing structural prediction tools could enhance their accuracy in modeling the folding and assembly of these complexes in three-dimensional space. Furthermore, this classification system can be used to validate predicted structures by comparing them to established interaction profiles, ensuring that the predicted conformations are both geometrically plausible and biologically relevant.

## CONCLUSIONS

ClaPNAC is the first classifier for amino acid–nucleotide/nucleoside interactions capable of annotating input 3D structures with pairwise contacts. Even very small changes in the input structure can be detected by ClaPNAC. The set of contact classes reported by ClaPNAC can be easily extended to include other types of structures or subtypes of spatial relationships between nucleotide and amino acid residues. In the future, ClaPNAC will also be expanded to enable the detection and classification of RNA–ligand interactions.

ClaPNAC represents a significant advancement in analyzing amino acid–nucleotide interactions by providing a comprehensive classification system that deepens our understanding of these crucial biological contacts. Its potential applications in 3D structure prediction and evaluation highlight its value as a powerful tool for researchers in structural biology.

## DATA AVAILABILITY

An implementation of ClaPNAC tool and the benchmarking data are available at https://genesilico.pl/software/stand-alone/clapnac and https://github.com/GrigorijNik/ClaPNAC (DOI: 10.5281/zenodo.15423175). An implementation of the contact extractor utility ContExt is available at https://github.com/febos/ContExt (DOI: 10.5281/zenodo.15374500)

## ACKNOWLEDGEMENTS

We would like to thank Angana Ray and Satyabrata Maiti for the preparation of dataset of RNP structures from PDB database.

## SUPPLEMENTARY INFORMATION

**Figure SF1.**
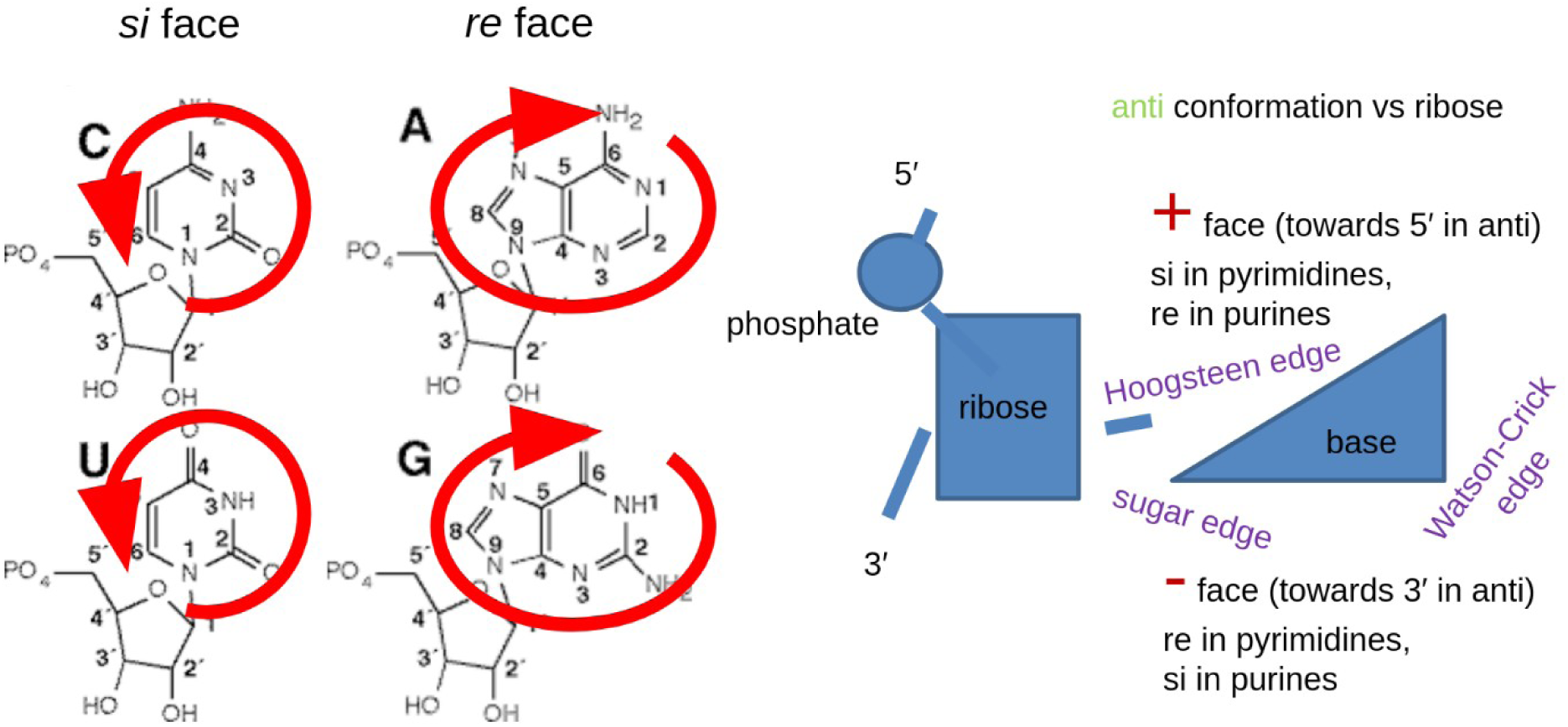
Interaction sites in ribonucleotides: base faces

**Figure SF2.**
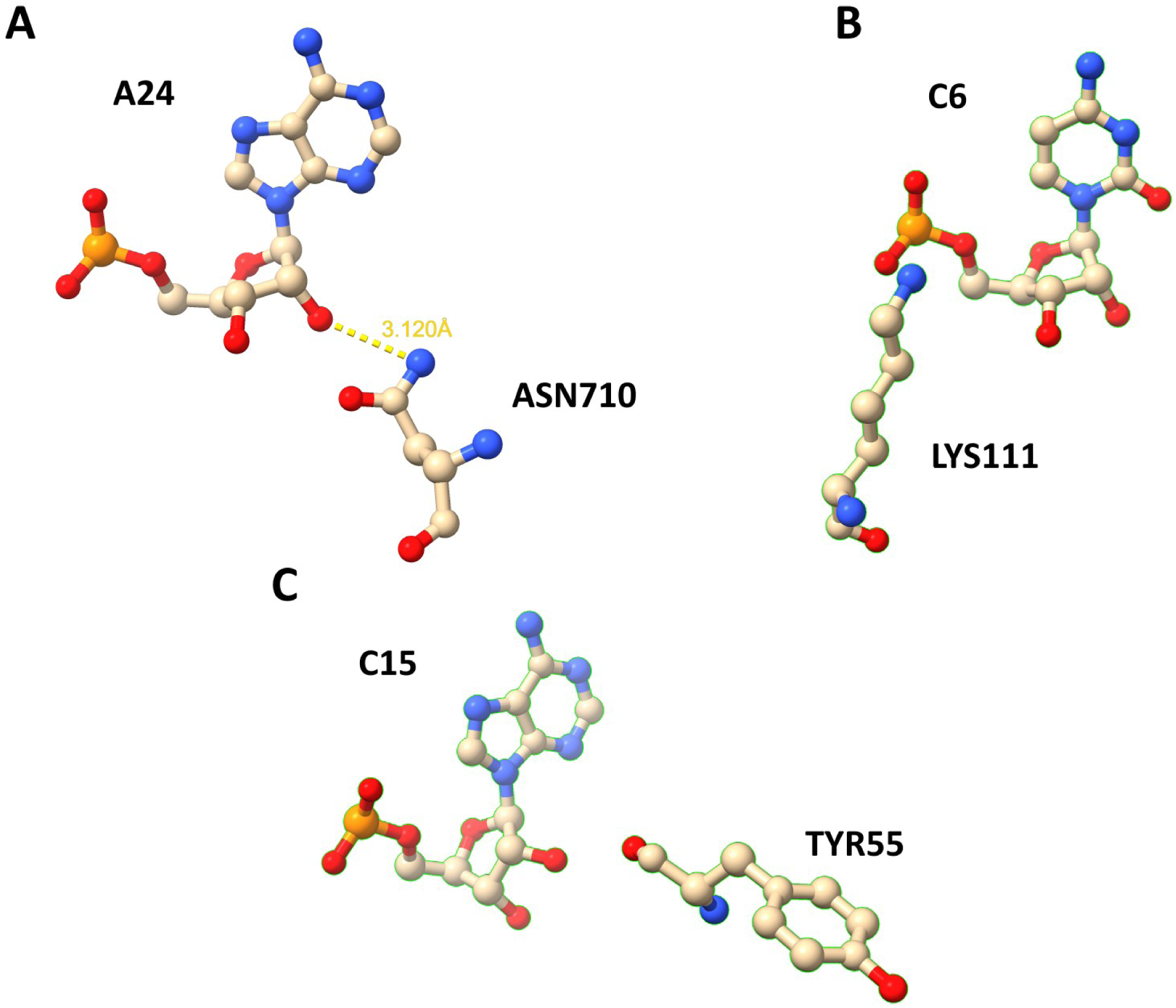
Examples of doublets that was not classified by ClaPNAC (A) Doublet from structure PDB id: 1FFY, (B) doublet from structure PDB id: 1JBS, (C) doublet from structure PDB id: 1JBT. Doublets A and B were classified by fingeRNAt and DSSR, but doublet C was classified only by fingeRNAt

---

**Table S1.** Number of nucleotide(N) - amino acid (AA) doublets.

|  | **ADE** | **CYT** | **GUA** | **URA** |
| --- | --- | --- | --- | --- |
| **ALA** | 769 | 356 | 768 | 1,012 |
| **ARG** | 10,203 | 10,518 | 12,135 | 9,325 |
| **ASN** | 2,179 | 1,574 | 2,069 | 2,733 |
| **ASP** | 1,353 | 710 | 2,075 | 1,332 |
| **CYS** | 86 | 22 | 65 | 132 |
| **GLN** | 1,970 | 1,501 | 2,443 | 2,145 |
| **GLU** | 1,038 | 834 | 1,808 | 1,177 |
| **GLY** | 2,065 | 1,558 | 2,371 | 1,695 |
| **HIS** | 2,026 | 2,110 | 2,406 | 1,753 |
| **ILE** | 451 | 386 | 504 | 508 |
| **LEU** | 617 | 523 | 775 | 768 |
| **LYS** | 5,306 | 4,780 | 6,637 | 4,463 |
| **MET** | 545 | 665 | 697 | 552 |
| **PHE** | 1,176 | 676 | 1,005 | 740 |
| **PRO** | 803 | 614 | 707 | 890 |
| **SER** | 3,896 | 2,365 | 3,238 | 2,020 |
| **THR** | 6,066 | 1,552 | 1,968 | 1,571 |
| **TRP** | 738 | 249 | 626 | 556 |
| **TYR** | 1,839 | 1,694 | 1,260 | 2,602 |
| **VAL** | 1,022 | 324 | 693 | 630 |
|  |  |  | **TOTAL** | 158,013 |

**Table S2.** Contacts in U5 snRNP State I, II, III, IV and U4/U6.U5 tri-snRNP.

| Nucleotide id | amino_acid id | score | interaction type | doublent | 8Q7V<br>(State I) | 8Q7Q<br>(State II) | 8Q7W<br>(State III) | 8Q7X<br>(State IV) | 6QW6 |
| --- | --- | --- | --- | --- | --- | --- | --- | --- | --- |
| 5-10 | A-217 | 1 | R | U-ARG | + | + | + | + | - |
| 5-11 | A-217 | 1 | R | U-ARG | + | + | + | + | + |
| 5-12 | A-222 | 1 | P | U-GLY | + | + | - | - | + |
| 5-12 | A-224 | 1 | P | U-THR | + | + | - | - | - |
| 5-13 | A-224 | 1 | P | C-THR | - | - | + | + | + |
| 5-14 | A-477 | 1 | P | U-LYS | + | + | + | + | - |
| 5-15 | A-474 | 1 | P | C-ARG | + | + | + | + | + |
| 5-17 | A-468 | 1 | + | U-LYS_Backbone | - | - | - | - | + |
| 5-17 | A-470 | 1 | P | U-ARG | - | - | - | - | + |
| 5-18 | A-468 | 1 | P | C-LYS | - | - | - | - | + |
| 5-18 | A-467 | 1 | H | C-GLN_Backbone | - | - | - | - | + |
| 5-20 | A-466 | 1 | P | G-ALA | + | + | + | + | - |
| 5-20 | A-463 | 1 | W | G-PRO_Backbone | + | + | + | + | - |
| 5-23 | A-461 | 1 | P | C-HIS | + | + | + | + | + |
| 5-23 | A-465 | 1 | H | C-LYS_Backbone | + | + | + | + | + |
| 5-26 | A-428 | 1 | P | A-LYS | + | + | + | + | + |
| 5-26 | A-635 | 1 | R | A-ARG | + | + | + | + | + |
| 5-26 | A-639 | 1 | R | A-PHE | + | + | + | + | + |
| 5-27 | A-428 | 1 | P | U-LYS | - | - | - | - | + |
| 5-27 | A-639 | 1 | R | U-PHE | + | + | + | + | + |
| 5-27 | A-457 | 1 | + | U-ASN_Backbone | + | + | + | + | + |
| 5-29 | A-597 | 1 | P | A-LYS | + | + | + | + | - |
| 5-29 | A-600 | 1 | P | A-ARG | - | - | - | - | + |
| 5-29 | A-644 | 1 | R | A-ILE | - | - | + | + | - |
| 5-30 | A-595 | 1 | P | A-LYS | + | + | - | - | + |
| 5-30 | A-597 | 1 | P | A-LYS | + | + | + | + | + |
| 5-35 | A-284 | 1 | R | U-ARG | - | - | - | - | + |
| 5-45 | A-603 | 1 | S | C-ARG | - | - | - | - | + |
| 5-45 | A-599 | 1 | S | C-MET | + | + | - | - | + |
| 5-45 | A-596 | 1 | - | C-TYR | + | + | - | - | - |
| 5-46 | A-603 | 1 | R | U-ARG | - | - | + | + | + |
| 5-47 | A-452 | 1 | P | A-LYS | - | - | + | + | + |
| 5-48 | A-452 | 1 | P | A-LYS | - | - | - | - | + |
| 5-49 | A-267 | 1 | P | A-LYS | + | + | + | + | + |
| 5-49 | A-460 | 1 | P | A-LYS | + | + | + | + | + |
| 5-51 | A-462 | 1 | P | A-ARG | + | + | + | + | - |
| 5-53 | A-465 | 1 | P | U-LYS | - | - | - | - | + |
| 5-55 | A-97 | 1 | R | C-HIS | - | - | - | - | + |
| 5-55 | A-642 | 1 | S | C-ARG | + | + | + | + | + |
| 5-56 | A-100 | 1 | R | C-LEU | - | - | - | - | + |
| 5-56 | A-420 | 1 | S | C-ARG | - | - | - | - | + |
| 5-57 | A-101 | 1 | P | G-LYS | + | + | + | + | + |
| 5-58 | A-134 | 1 | P | U-TRP | + | + | + | + | + |
| 5-59 | A-469 | 1 | H | G-LYS | + | + | + | + | + |
| 5-60 | A-469 | 1 | H | G-LYS | + | + | - | - | + |
| 5-64 | A-50 | 1 | P | G-LYS | - | - | - | - | + |

**Table S3.** Doublets with nucleotide R663 classified by ClaPNAC with SF = 1 from 3MUM, 3MUR, 3IRW.

| nucleotide id | amino_acid id | Score | interaction type | doublet | 3IR W | 3MU M | 3MU R |
| --- | --- | --- | --- | --- | --- | --- | --- |
| R-61 | P-22 | 1 | P | A-LYS | - | - | + |
| R-62 | P-22 | 1 | P | A-LYS | + | - | + |
| R-62 | P-47 | 1 | P | A-ARG | + | + | - |
| R-63 | P-20 | 1 | P | A-LYS | + | - | + |
| R-64 | P-20 | 1 | P | C-LYS | - | - | + |
| R-660 | P-52 | 1 | W | A-ARG | + | + | + |
| R-661 | P-52 | 1 | S | U-ARG | + | - | - |
| R-661 | P-19 | 1 | W | U-GLU | + | + | + |
| R-662 | P-16 | 1 | W | U-ASN_Backbone | + | + | + |
| R-662 | P-80 | 1 | W | U-LYS | + | + | + |
| R-663 | P-50 | 1 | R | G-LYS | - | - | + |
| R-663 | P-54 | 1 | + | G-GLN | + | + | + |
| R-663 | P-15 | 1 | + | G-ASN | + | + | + |
| R-663 | P-52 | 1 | S | G-ARG_Backbone | + | + | - |
| R-663 | P-53 | 1 | - | G-GLY | + | + | + |
| R-663 | P-19 | 1 | W | G-GLY | - | + | + |
| R-664 | P-13 | 1 | - | C-TYR | + | + | + |
| R-664 | P-87 | 1 | W | C-ALA | + | - | - |
| R-664 | P-88 | 1 | + | C-LYS | + | - | + |
| R-665 | P-51 | 1 | R | A-MET | + | + | + |
| R-665 | P-56 | 1 | - | A-PHE | + | - | + |
| R-665 | P-91 | 1 | W | A-SER | + | - | + |
| R-666 | P-92 | 1 | + | C-ASP | + | - | - |
| R-666 | P-92 | 1 | W | C-ASP | + | + | + |
| R-666 | P-91 | 1 | W | C-SER | + | + | + |
| R-669 | P-50 | 1 | + | C-LYS | - | + | - |
| R-75 | P-52 | 1 | W | G-ARG | - | - | + |

**Table S4.** Frequences of the base/backbone interactions, interaction classes, and the nucleotides/nucleosides and amino acids for PF00076 and PF04059 Pfam families, RNA-Protein and DNA-Protein datasets.

|  | PF00076 | PF04059 | RNA-Protein | DNA-Protein |
| --- | --- | --- | --- | --- |
|  | Base/backbone interaction frequencies |  |  |  |
| **Backbone** | 0.7118 | 0.2766 | 0.7093 | 0.7303 |
| **Base** | 0.2882 | 0.7234 | 0.2907 | 0.2697 |
|  | Interactions classes frequencies |  |  |  |
| **P** | 0.4347 | 0.1489 | 0.3794 | 0.5333 |
| **R** | 0.2772 | 0.1277 | 0.3299 | 0.197 |
| **W** | 0.0766 | 0.3191 | 0.0845 | 0.0045 |
| **H** | 0.0588 | 0.0426 | 0.0433 | 0.1818 |
| **S** | 0.0351 | 0.0693 | 0.0474 | 0.0167 |
| **+** | 0.0654 | 0.1915 | 0.0464 | 0.0439 |
| **-** | 0.0522 | 0.1009 | 0.0691 | 0.0228 |
|  | Nucleotide/nucleoside frequencies |  |  |  |
| **A** | 0.2039 | 0.0612 | 0.1835 | 0.2061 |
| **C** | 0.2112 | 0.0426 | 0.2412 | 0.2621 |
| **G** | 0.3132 | 0.1277 | 0.2165 | 0.2697 |
| **U/T** | 0.2717 | 0.7685 | 0.3588 | 0.2621 |
|  | Amino acid frquences |  |  |  |
| **PHE** | 0.0293 | 0.1277 | 0.0412 | 0.0106 |
| **LYS** | 0.2604 | 0.1064 | 0.1691 | 0.2061 |
| **ARG** | 0.2425 | 0.0851 | 0.1732 | 0.1788 |
| **ILE** | 0.014 | 0.2128 | 0.0258 | 0.0212 |
| **LEU** | 0.0172 | 0 | 0.0186 | 0.0061 |
| **TYR** | 0.0377 | 0.1489 | 0.1227 | 0.0788 |
| **THR** | 0.0502 | 0 | 0.1227 | 0.0788 |
| **HIS** | 0.0457 | 0 | 0.0404 | 0.05 |
| **TRP** | 0.0232 | 0 | 0.0155 | 0.0515 |
| **ASN** | 0.0226 | 0.0426 | 0.0206 | 0.0484 |
| **GLY** | 0.0702 | 0.0213 | 0.0887 | 0.0712 |
| **SER** | 0.0616 | 0.0426 | 0.0691 | 0.0803 |
| **ALA** | 0.0293 | 0.0851 | 0.0206 | 0.0409 |
| **ASP** | 0.013 | 0 | 0.0237 | 0.0393 |
| **PRO** | 0.019 | 0 | 0.0237 | 0.0152 |
| **VAL** | 0.02 | 0 | 0.0319 | 0.0106 |
| **GLN** | 0.0199 | 0 | 0.0227 | 0.0394 |
| **MET** | 0.0095 | 0.0426 | 0.0103 | 0.0076 |
| **GLU** | 0.0094 | 0 | 0.0175 | 0.0061 |
| **CYS** | 0.002 | 0.0213 | 0.0021 | 0 |

